# The evolution of parasite host range in genetically diverse host populations

**DOI:** 10.1101/653675

**Authors:** Amanda K Gibson, Helena S Baffoe-Bonnie, McKenna J Penley, Julie Lin, Raythe Owens, Arooj Khalid, Levi T. Morran

## Abstract

Parasites vary enormously in their host range. Why are some parasites specialists and others generalists? We tested the hypothesis that genetically diverse host populations select for parasites with broader host ranges than genetically homogeneous host populations. To do so, we selected for increased killing ability of the bacterial parasite *Serratia marcescens* in populations of the host *Caenorhabditis elegans* that were either diverse (50% mix of genotypes N2 and LTM1) or homogenous (100% N2 or LTM1). We found mixed support for the hypothesis. After 20 generations of selection, parasites selected in diverse host populations maintained a broad host range, as shown by the retention of high killing ability of a novel host genotype. Parasites selected in diverse populations killed N2 hosts equally well as parasites selected in homogenous populations of N2. However, N2-selected parasites lost killing ability against the novel host, consistent with the evolution of narrow host ranges in homogenous environments. In contrast, parasites selected in homogenous LTM1 populations did not specialize: they did not increase in their killing ability on LTM1 and did not lose killing ability against the novel host. Our results argue that the evolution of host range depends upon both the identity and diversity of hosts that a parasite encounters.

## Introduction

Closely-related parasites may vary in the number of host genotypes (Carius et al. 2001; Thrall et al. 2001; Poullain et al. 2008), species (Desdevises et al. 2002; Poulin and Mouillot 2003; Krasnov et al. 2004), or even families (Ross et al. 2013) they can attack. Why does host range vary so much? Here, we use experimental evolution to test the hypothesis that the variance of the host environment drives the evolution of parasite host range, such that genetically diverse host populations favor generalist parasites and genetically homogeneous host populations favor specialist parasites.

This hypothesis derives from theory on the evolution of niche width. This body of theory argues that heterogeneous environments select for large niche widths (i.e. generalists), while homogeneous environments select for narrow niche widths (i.e. specialists) (Levins 1962; Pianka 1966; Via and Lande 1985; Lynch and Gabriel 1987; Futuyma and Moreno 1988). Specialization in a homogeneous environment is thought to arise from either antagonistic pleiotropy, where mutations that increase performance in the focal environment reduce performance in alternate environments (Rausher 1984; Jaenike 1990; Via 1990), or mutation accumulation, where populations accumulate mutations that are neutral in the focal environment and deleterious in alternate environments (Fry 1996; Whitlock 1996). The probability of fixation of such mutations declines if individuals have a high probability of encountering multiple environments, due to either temporal or fine-scale spatial heterogeneity. Experimental evolution studies of free-living systems support the maintenance of generalists under abiotic heterogeneity, notably under temporal heterogeneity (Bennett et al. 1992; Reboud and Bell 1997) (rev. in Kassen 2002).

This body of theory has been extended to the evolution of host range in parasites. Substantial evidence now exists for the evolution of host specialization under temporal homogeneity: during serial passage, many parasites adapt to infect individual host species or genotypes and simultaneously decline in their ability to infect alternate hosts (Cunfer 1984; Fry 1990; Ebert 1998). In contrast, tests of the associated prediction, that diverse host environments select for generalist parasites, are few in number and limited to viral systems. A common approach involves creating temporal host heterogeneity by alternating a viral lineage between cell lines derived from different host species (Novella et al. 1999; Weaver et al. 1999; Coffey and Vignuzzi 2011) (though see Bedhomme et al. 2011). Turner et al. (2010) adopted this approach, finding mixed support for the idea that diverse host communities select for parasites that can infect a broad range of host species: after alternation between human and canine cells, populations of vesicular stomatitis virus populations showed higher performance on cell lines of novel host species relative to viral populations serially passaged on human cells. In contrast, viral populations serially passaged on canine cells showed no average reduction in performance relative to viral populations selected under temporal alternation. Studies have not explicitly tested if spatial variation in hosts favor parasites with broad host ranges.

We build on these studies by directly testing the hypothesis that genetically diverse host populations select for parasites with a broader range of host genotypes than genetically homogeneous host populations. To test this hypothesis, we used experimental evolution to select on populations of the bacterial parasite *Serratia marcescens* for increased performance in populations of the nematode host *Caenorhabditis elegans* that varied in their diversity. Some host populations were genetically diverse (an even mix of two genotypes), creating fine-scale spatial variation in the host environment. Others were genetically homogeneous (one of two possible genotypes), creating spatial homogeneity. We then compared the breadth of host ranges across treatments by evaluating the performance of evolved parasite lineages on a novel host genotype. We predicted that 1) parasites selected in diverse host populations would maintain or increase in their ability to kill a novel host genotype, consistent with selection for a broad host range, and 2) parasites selected in homogeneous populations would show reduced killing ability of the novel host, consistent with specialization. We found mixed support for these predictions.

## Materials and Methods

### Host and parasite genotypes

For experimental evolution, we used two genotypes of the nematode *Caenorhabditis elegans:* N2 and LTM1. Slowinksi et al. (2016) described the origins of the LTM1 line, which is a single genotype derived from ethylmethane sulfonate mutagenesis of the CB4856 genotype. We selected these two host genotypes for experimental evolution because 1) N2 and CB4856 are among the most genetically divergent genotypes within *C. elegans* (Barrière and Félix 2005), and 2) preliminary assays demonstrated that the parasite *Serratia marcescens* is equally virulent to N2 and LTM1.

For assays of parasite virulence, we also included the host genotype JU1395. JU1395 is roughly equally genetically divergent from N2 and LTM1(Andersen et al. 2012). Hence we limited the potential that genetic proximity alone would generate differences between parasites adapted to N2 vs. LTM1 in their virulence against JU1395. Assays of parasite performance against JU1395 allowed us to compare the host range of evolved parasite lineages. We subsequently refer to JU1395 as a novel host genotype, because parasite lineages never encountered this host genotype during experimental evolution. We refer to N2 and LTM1 as sympatric host genotypes, because parasite lineages encountered one or both of these host genotypes during experimental evolution.

We initiated replicate parasite lineages from Sm2170, a genotype of the bacterial parasite *Serratia marcescens*. Sm2170 is known to be highly virulent towards *C. elegans* hosts (Schulenburg and Ewbank 2004). The interaction of *C. elegans* and Sm2170 is a novel host-parasite interaction constructed in the lab: there is no evidence that *C. elegans* encounters this particular strain of *S. marcescens* in the wild, and Sm2170 had not previously been experimentally evolved with *C. elegans*. Hosts acquire infection while feeding.

### Parasite selection treatments

We established four treatments, each with six replicate parasite lineages (Fig. 1). In three of these treatments, we subjected replicate parasite lineages to direct selection for increased virulence against host populations that differed in their level of genetic diversity. In the first two treatments, parasites were selected to kill hosts in homogeneous host populations. These host populations comprised either 100% N2 hosts or 100% LTM1 hosts. In the third treatment, parasites were selected to kill hosts in populations that were genetically diverse. These populations were 50% N2 and 50% LTM1 hosts. We assumed that parasites have no ability for host choice, such that parasites passaged with genetically diverse host populations had an equal probability of encountering an N2 or LTM1 host each generation of selection.

**Figure 1:**
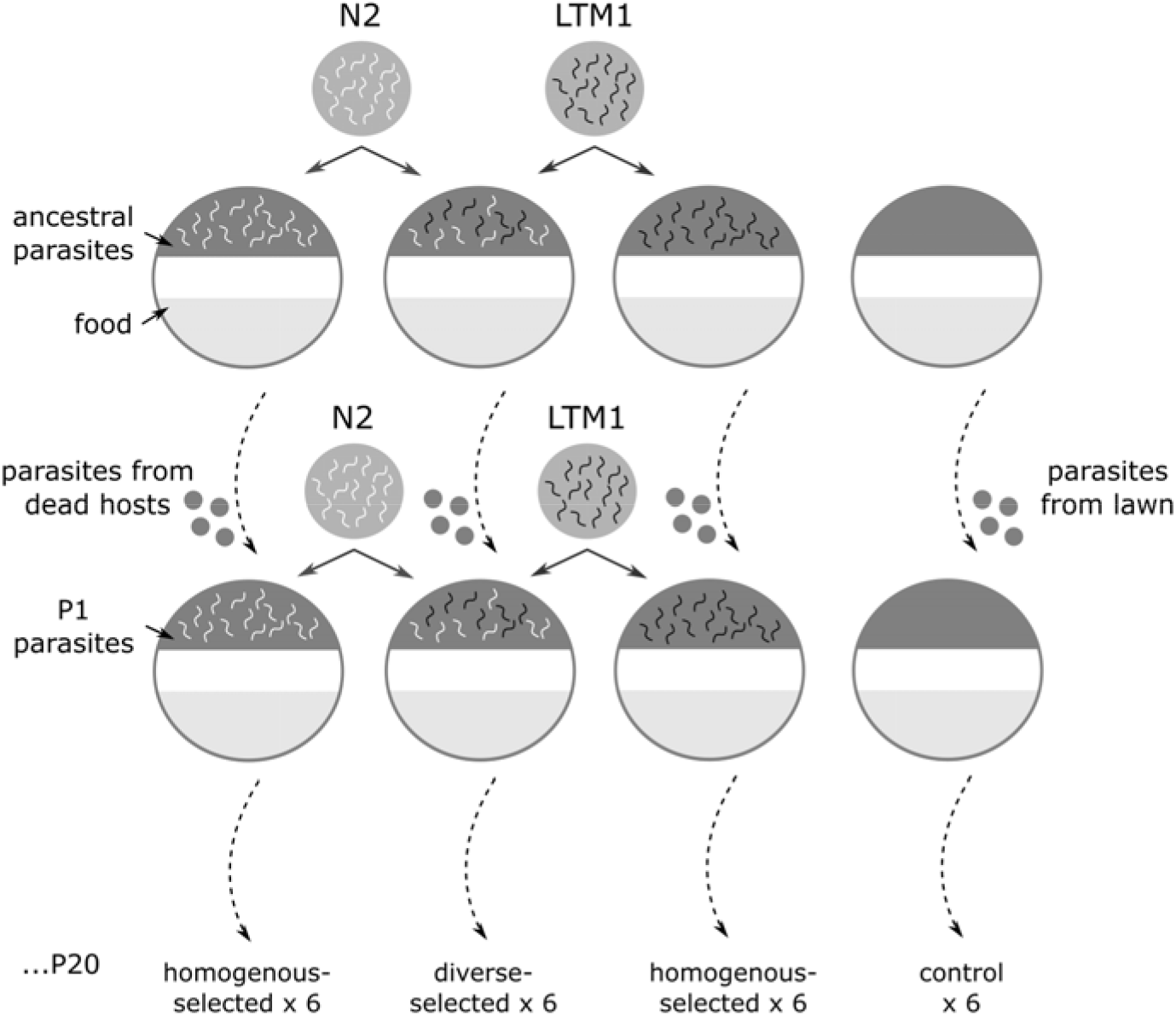
Experimental evolution scheme. We initiated experimental evolution by adding 500 *C. elegans* hosts to *Serratia* selection plates seeded with a lawn of Sm2170, the ancestral parasite genotype (dark lawn on upper portion of plates). For homogeneous selection, we added 100% N2 (left, white) or 100% LTM1 (right, black) hosts. For diverse selection, we added 50% N2 and 50% LTM1 hosts. We then selected for virulent parasites by extracting parasite colonies from hosts that died rapidly, within 24 hours. We used this passage of parasites (shown here as P1, second row) to seed lawns on *Serratia* selection plates, to which the same genotype(s) of hosts were added to commence the second round of selection. For the control treatment, we did not add any hosts and passaged parasite colonies directly from the lawn. We continued these selection regimes for a total of 20 passages. Each of the four treatments was replicated six times, for a total of 24 independent parasite lineages.

We did not allow for host evolution during experimental evolution. Hence, each passage, parasites were re-exposed to host populations of the same make-up as the prior generation. We limited host evolution by maintaining stock populations of N2 and LTM1 at 15°C. Every few weeks, we refreshed these stocks by thawing hosts archived at −80°C. Our experimental treatments therefore limited temporal host heterogeneity in order to contrast spatial host heterogeneity with homogeneity.

The fourth treatment was the control treatment, where we did not directly select for increased virulence. We designed this treatment to serve as the baseline against which to measure evolutionary change in the prior three treatments. In this treatment, we passaged bacteria without hosts. In doing so, we subjected bacterial populations to genetic drift and to the non-focal selection pressures of the experiment in the absence of selection for increased virulence.

### Experimental evolution design

Selection was performed using *Serratia* selection plates, as in Morran et al. (2009) (Fig. 1). Under this design, we seeded 100 mm petri dishes of Nematode Growth Media (NGM-lite, United States Biological) with 35 uL of bacterial parasites (*Serratia marcescens*) on one side of the plate and 35 uL of food (*Escherichia coli*, strain OP50) on the other. Adding nematodes to the *Serratia* lawn forced interaction between hosts and parasites. Hosts could then migrate towards the lawn of food. We used this particular design in order to maintain the conditions of prior evolution experiments (Morran et al. 2009; Morran et al. 2011; Slowinski et al. 2016) and thereby facilitate comparison with their results.

To initiate experimental evolution, we harvested large numbers of L4 larvae of N2 and LTM1 hosts. We established host populations that were 100% N2, 100% LTM1, or 50% N2:50% LTM1 hosts by mixing the appropriate volumes of larvae of each host genotype. All initial *Serratia* selection plates were seeded with the same culture of Sm2170. In order to establish six replicate parasite lineages per treatment, we deposited ~500 L4 larvae of the appropriate host population onto the Sm2170 lawns of six different *Serratia* selection plates. For the control treatment, we did not add any larvae to the Sm2170 lawns. This resulted in a total of 24 plates representing 24 independent parasite lineages, six per each of four treatments.

We maintained these plates for 24 hours at 20°C. We then selected the most virulent parasites by isolating and transferring those that killed hosts rapidly, within 24 hours. To accomplish this, we picked 20-30 dead hosts from the Sm2170 lawn of each plate. We removed external bacteria from these hosts by repeated rinsing, then crushed the hosts to extract the internal bacteria that had killed them (Morran et al. 2011). We grew these bacteria on NGM-lite plates at room temperature (~22°C) for 48 hours, then maintained them at 4°C for 48 hours. We then randomly selected 40 colonies from these plates and grew them at 28°C overnight in 5 mL of LB media. These liquid cultures were used to produce the next round of *Serratia* selection plates, to which we added the same host population encountered by the parasite lineage in the prior passage.

For the control treatment, we collected ~30 samples of free-living bacteria directly from the lawn of *Serratia* in order to mimic the sample sizes obtained in the other treatments. We otherwise treated these populations in the same manner as the host-associated lineages. We repeated this passaging scheme for a total of 20 passages, at which point we froze liquid cultures of parasite lineages at −80°C.

### Survival assays of parasite virulence

We measured parasite virulence as the mortality rate of a host genotype after 48 hours of exposure to a parasite lineage. In setting up the assays, we replicated the experimental passaging scheme. For each host genotype tested, we added a fixed volume of L4 larvae (100% focal host genotype) to multiple replicate *Serratia* selection plates of all 24 parasite lineages. We determined the mean number of L4 larvae added to *Serratia* selection plates by adding this same volume to 10 standard plates seeded with OP50 and counting the number of hosts after 24 hours. We maintained *Serratia* selection plates at 20°C for 48 hours, then counted the number of live worms that had migrated out of the *Serratia* lawn. The mortality rate was obtained from the survival rate, which we calculated as the number of live hosts divided by the mean number added.

For the N2 genotype, we added 494 ± 26 hosts (mean ± standard error of the mean) per *Serratia* selection plates. Each parasite lineage was replicated four times, for a total of 24 experimental replicates per selection treatment. For the LTM1 genotype, we added 498 ± 25 hosts. Each parasite lineage was also replicated four times. For our novel genotype, JU1395, we added 270 ± 12 hosts. Each parasite lineage was replicated eight times, for a total of 48 experimental replicates per selection treatment.

### Statistical Analyses

All statistical analyses were performed in R (ver. 3.5.3; R Core Team 2013). We conducted three separate analyses, one for each host genotype tested in the survival assays, in order to compare the virulence of parasite lineages from different experimental treatments towards a given host genotype. Statistical analyses with the N2 and LTM1 genotypes served to evaluate adaptation of parasite lineages to their sympatric host genotypes. The statistical analysis with the JU1395 genotype served to evaluate the host range of selected parasite lineages, by testing their ability to kill a novel host genotype.

We began with survival assay data for N2, one of the sympatric host genotypes. We fit a Poisson regression with experimental evolution treatment (control, homogeneous N2, homogeneous LTM1, diverse) as a predictor of the number of live worms in an experimental replicate. We included parasite lineage (1-6) as a random effect. We found evidence of significant overdispersion (variance inflation factor, *ĉ*=19.98)(Venables and Ripley 2002), so we re-fit the model as a negative binomial regression with the glmer.nb function in the lme4 package (Bates et al. 2015). A likelihood ratio test indicated a substantially better fit with the negative binomial regression relative to the Poisson regression (Likelihood-ratio test: χ^2^=1319.2, df=1, p<0.001). We applied this same modeling approach for the LTM1 and JU1395 genotypes. In both cases, we found evidence of overdispersion (LTM1, *ĉ*=21.65; JU1395, *ĉ*=11.63) and a better fit to our data with a negative binomial regression (LTM1, χ^2^=1504.5, df=1, p<0.001; JU1395, χ^2^=1448.1, df=1, p<0.001).

We then evaluated treatment as a predictor of variation in the number of surviving hosts by using likelihood ratio tests to compare models with and without the treatment factor. For models in which treatment was a significant predictor of variation in survival, we examined model coefficients to compare between treatments. In analysis of sympatric host genotypes, we tested the prediction that parasite lineages evolved increased virulence against hosts with which they were passaged during experimental evolution. In analysis of the novel host genotype, we tested the prediction that parasite lineages selected in diverse host populations would have higher virulence against a novel host than parasite lineages selected in homogeneous host populations, consistent with a larger host range for parasites selected in diverse host populations.

Lastly, we conducted a post-hoc analysis, based on observation of the data, to test if parasite lineages selected in diverse host populations varied less in their virulence against a novel host than parasite lineages selected in homogeneous host populations. To test this prediction, we calculated the coefficient of variation in virulence (both number of surviving hosts and mortality rate) against JU1395 across the six independent parasite lineages per treatment. We calculated 95% confidence intervals for the coefficient of variation by bootstrapping the JU1395 data set 10,000 times. Specifically, we re-sampled the experimental replicates per parasite lineage eight times with replacement and re-calculated the coefficient of variation for each treatment.

## Results

### Adaptation to sympatric host genotypes

We first evaluated the virulence of experimentally evolved parasites when paired with their sympatric hosts, N2 and LTM1. We predicted an increase in virulence when parasite lineages were paired with the host genotypes on which they were selected.

The mortality rate of N2 was 82.3% when paired with the ancestral parasite genotype. This closely matched the mortality rate of N2 when paired with control parasite lineages after 20 experimental passages (82.6 ± 0.8% SEM) (Table S1). When paired with experimentally evolved parasites, the mortality rate of N2 varied with parasite selection treatment (Table 1A, Fig. 2A). Consistent with our prediction, parasite lineages selected to kill hosts in populations that were diverse or homogeneous for N2 showed increased ability to kill N2 hosts relative to control parasites (Generalized linear mixed model, number of surviving worms relative to control: 50%: coefficient = −0.315, *z* = −2.287, *p* = 0.022; 100% N2: coefficient = −0.514, *z* = −3.677, *p* < 0.001) and parasites selected to kill homogeneous LTM1 populations (50%: coefficient = 0.383, *z* = 2.755, *p* = 0.006; 100% N2: coefficient = 0.582, *z* = 4.232, *p* < 0.001). Specifically, the mortality rate of N2 was 87.3±0.7% and 88.4±1.4% for parasites selected in diverse or homogeneous N2 populations, respectively. Therefore, in relation to control parasites, survival of N2 hosts declined by approximately a third when paired with parasites selected to kill N2 (homogeneous N2: 33% decline; diverse: 27% decline). Parasites selected in diverse or homogeneous N2 populations did not differ from one another in their killing ability (coefficient = 0.198, *z* = 1.417, *p* = 0.157). Parasite lineages selected to kill hosts in populations that were homogeneous for LTM1 showed equivalent ability to kill N2 as control parasites (79.7 ± 2.1%, coefficient = 0.068, *z* = 0.494, *p* = 0.621).

**Table 1:** Experimental evolution treatment as a predictor of variation in parasite virulence on sympatric and novel host genotypes

|  | df | *D* | *p* |
| --- | --- | --- | --- |
| *On sympatric hosts* |  |  |  |
| A. N2 | 3 | 22.93 | <0.001 |
| B. LTM1 | 3 | 3.73 | 0.292 |
| *On novel host* |  |  |  |
| C. JU1395 | 3 | 30.01 | <0.001 |

Results of three separate generalized linear mixed models in which we fit parasite experimental evolution treatment (homogeneous N2, homogeneous LTM1, diverse, or control) as a predictor of the number of host individuals that survived parasite exposure. The three models correspond to three separate killing assays, one for each host genotype tested (A - N2, B - LTM1, and C - JU1395). We included parasite lineage (six independent lineages per experimental evolution treatment) as a random effect. We show the results of likelihood ratio (*D*) tests of models with and without treatment as a predictor.

**Figure 2:**
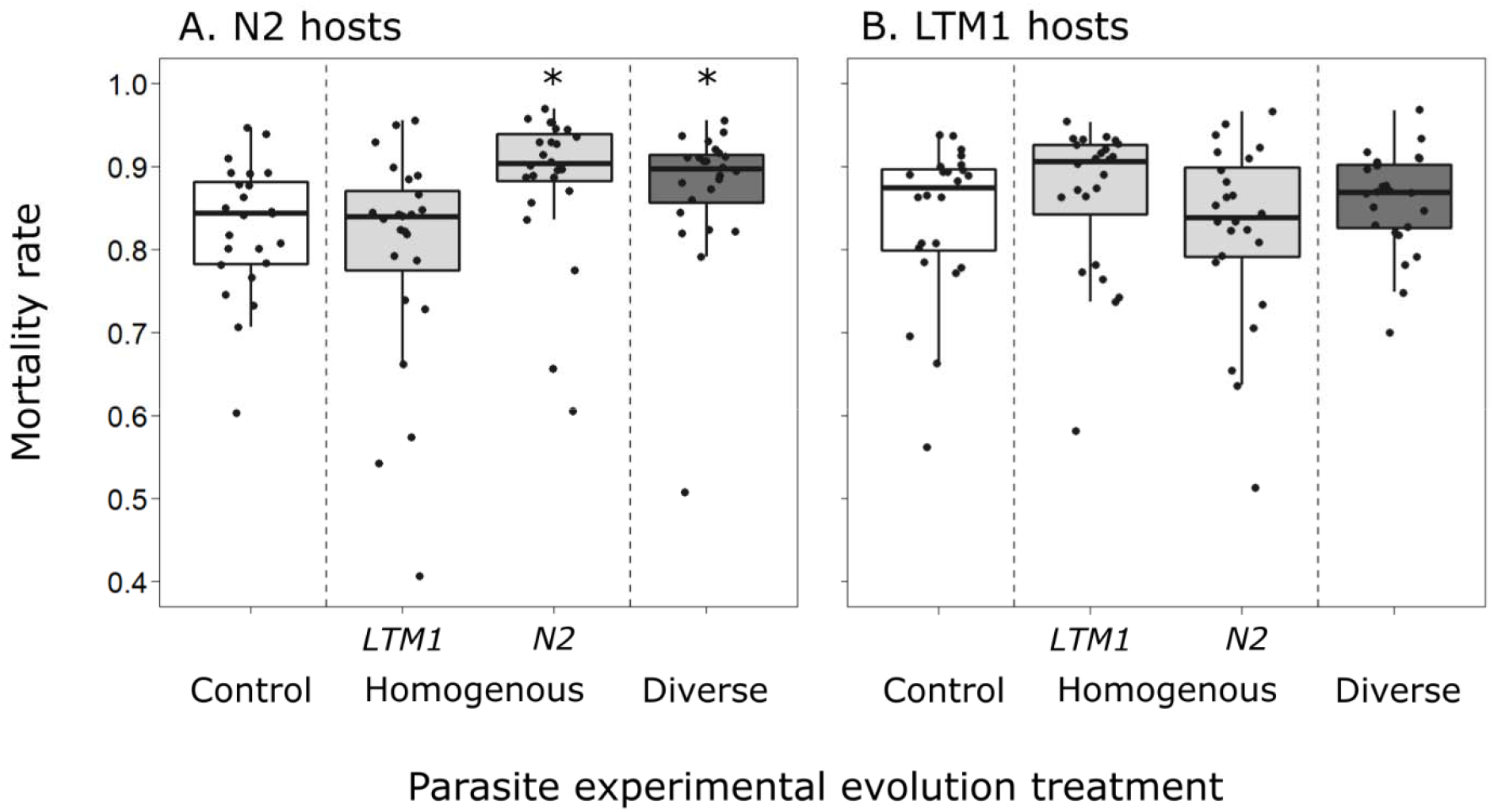
Virulence of experimentally evolved parasites on their sympatric host genotypes. The parasite *Serratia marcescens* was selected to kill *C. elegans* hosts in host populations that were homogeneous (100% LTM1; 100% N2) or diverse (50% LTM1: 50% N2). After 20 passages, we then tested evolved parasite lineages for their ability to kill N2 and LTM1 hosts. We compared the mortality rate of parasites against these hosts to that of control parasites, which were not selected to kill hosts and hence reflect baseline killing ability. (A) Parasites selected to kill hosts in populations that were diverse or homogeneous for N2 evolved an increased ability to kill N2 hosts, relative to control parasites or parasites selected to kill hosts in populations that were homogeneous for LTM1. (B) In contrast, parasites from different experimental evolution treatments did not differ in their ability to kill LTM1. Each box summarizes the results of 24 experimental replicates (4 replicates for each of 6 parasite lineages per treatment). Each point shows the mortality rate in a single experimental replicate, with 494 ± 26 (N2) or 498 ± 25 (LTM1) hosts tested per replicate.

When paired with the ancestral parasite genotype, the mortality rate of LTM1 was 82.2%, identical to that of N2 hosts. This mortality rate was slightly lower than the mortality rate of LTM1 when paired with control parasite lineages after 20 experimental passages (83.8 ± 0.5% SEM) (Table S2). Counter to our prediction, and in contrast to the results obtained for the N2 host genotype, the mortality rate of LTM1 did not vary across selection treatments for experimentally evolved parasite lineages (Table 1B, Fig. 2B). The changes in virulence qualitatively matched those observed with N2: relative to control parasites, the mortality rate of LTM1 was slightly higher when paired with parasites selected in populations that were diverse or homogeneous for LTM1 (85.8 ± 0.6%, 86.5 ± 0.8%). We likewise observed no increase from control parasites in the mortality rate of LTM1 when paired with parasite lineages selected to kill hosts in populations that were homogeneous N2 (82.3 ± 1.5%).

### Adaptation to a novel host genotype

We then evaluated the virulence of experimentally evolved parasites when paired with a novel host genotype, JU1395. We initially predicted 1) an increase or maintenance of virulence against the novel host genotype for parasites selected in diverse host populations and 2) a decrease in virulence against the novel host genotype for parasites selected in homogeneous host populations. Our results on adaptation to sympatric host genotypes subsequently suggested that support for these predictions would be strongest in comparisons of parasites selected on diverse populations vs. populations homogeneous for N2.

The mortality rate of JU1395 was 95.6% when paired with the ancestral parasite genotype. This mortality rate was slightly higher than the mortality rate of JU1395 when paired with control parasite lineages after 20 experimental passages (93.6 ± 0.3% SEM) (Table S3). When paired with experimentally evolved parasites, the mortality rate of JU1395 varied with parasite selection treatment (Table 1C, Fig. 3).

**Figure 3:**
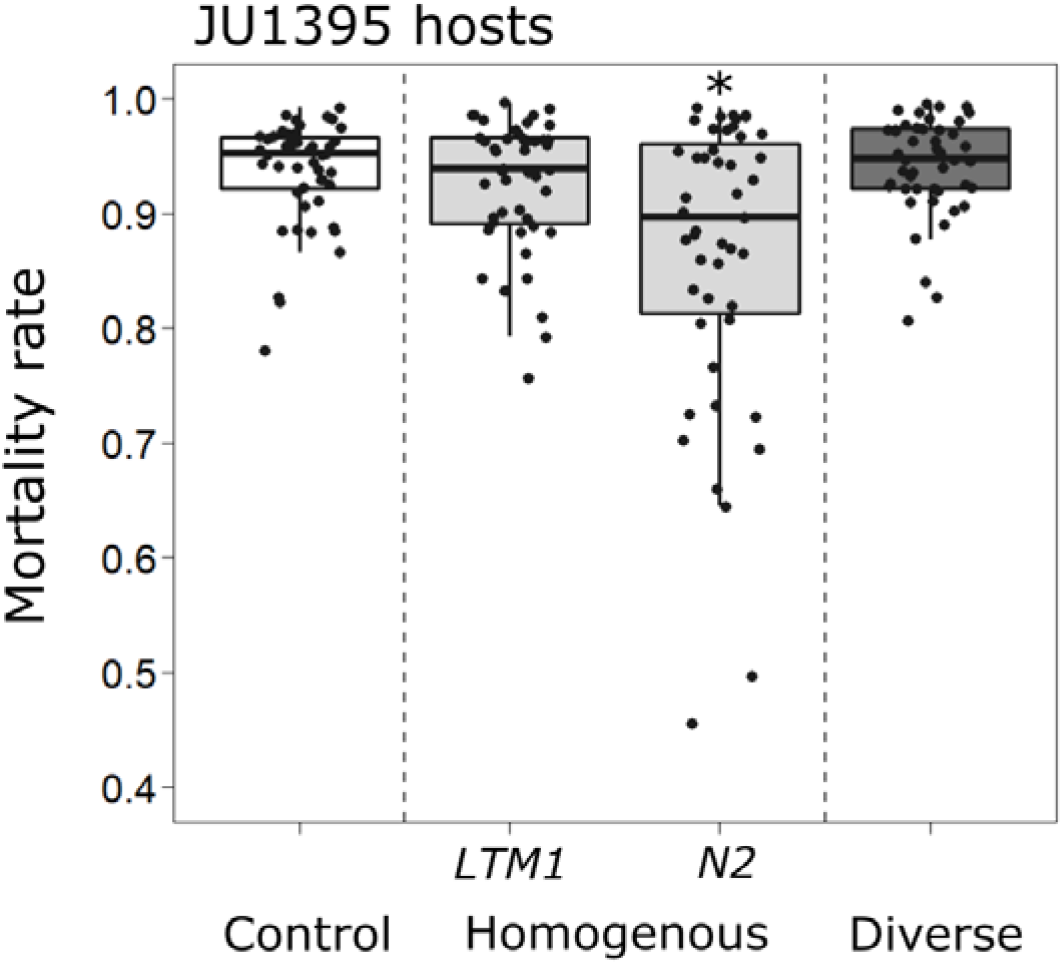
Virulence of experimentally evolved parasites on a novel host genotype. Here, we tested evolved parasite lineages for their ability to kill a novel host genotype, JU1395. Parasites selected to kill hosts in populations that were homogeneous for N2 (100% N2) lost their ability to kill the novel host (reduced mortality rate), consistent with specialization on the N2 genotype. In contrast, parasites selected to kill hosts in populations that were diverse (50% N2: 50% LTM1) maintained their ability to kill the novel host (equivalent mortality rate), consistent with the maintenance of generalism. Parasites selected to kill hosts in populations were homogeneous for LTM1 (100% LTM1) also showed no change in killing ability of the novel host, relative to control parasites. Each box summarizes the results of 48 experimental replicates (8 replicates for each of 6 parasite lineages per treatment). Each point shows the mortality rate in a single experimental replicate, with 270 ± 12 hosts tested per replicate.

We found support for our first prediction, that parasites selected in diverse populations would maintain killing ability against a novel host genotype. Parasite lineages selected to kill hosts in populations that were diverse showed an equivalent ability to kill JU1395 as control parasites (94.1 ± 0.3%; coefficient = −0.12856, *z* = −0.836, *p* = 0.403).

We found partial support for our second prediction that parasites selected in homogeneous host populations would lose killing ability against a novel host. Parasite lineages selected to kill hosts in populations that were homogeneous for N2 showed reduced ability to kill JU1395 hosts relative to control parasites (coefficient = 0.632, *z* = 4.075, *p* < 0.001) and to parasites selected to kill hosts in diverse populations (coefficient = 0.761, *z* = 4.962, *p* < 0.001). Specifically, the mortality rate of JU1395 was 86.1 ± 1.8% when paired with parasites selected to kill hosts in populations that were homogeneous N2. Therefore, in relation to control or diverse-selected parasites, survival of novel hosts was more than two-fold greater with parasites selected in homogeneous N2 populations (vs. control: 2.19-fold greater; vs. 50%: 2.35-fold greater). In contrast, the mortality rate of JU1395 was 92.5 ± 0.7% when paired with parasite lineages selected to kill hosts in populations that were homogeneous LTM1. This did not differ significantly from mortality rates of JU1395 hosts when paired with control parasites (coefficient = 0.093, *z* = 0.606, *p* = 0.544) or diverse-selected parasites (coefficient = 0.221, *z* = 1.442, *p* = 0.149). While counter to our prediction, this latter result corresponds to our finding that parasites failed to evolve significantly increased virulence against LTM1 (Table 1B, Fig. 2B).

Consistent with our findings above, parasites selected in homogeneous N2 populations showed the greatest between-lineage variation in performance on the novel host genotype. Control parasites and diverse-selected parasites showed equivalent between-lineage variation in their ability to kill the novel host, both in terms of number of survivors (coefficients of variation: 0.299, 95% CI [0.155,0.582] v. 0.312 [0.219,0.552], respectively) and mortality rate (0.020 [0.010,0.042] v. 0.020 [0.014,0.035]) (Table S4). For parasites selected in populations that were homogeneous N2, between-lineage variation was substantially higher (number of survivors: 0.780 [0.622,0.965]; mortality rate: 0.126 [0.092, 0.169]). This variation arose from the fact that virulence against JU1395 was very low for some lineages in this treatment (mortality rates: 69.5 ± 4.3%, 75.0 ± 4.8%) and high for others (92.9 ± 1.6%, 93.1 ± 2.5%, 93.4 ± 2.0%, 92.6 ± 1.8%). For parasites selected in populations that were homogeneous LTM1, between-lineage variation was somewhat elevated (number of survivors: 0.565 [0.413,0.779]; mortality rate: 0.046 [0.032, 0.065]), though not to the same extent as observed for parasites selected in homogeneous N2 populations.

## Discussion

We set out to test the hypothesis that parasite host range evolves to match the variance of the host environment. Specifically, we predicted that genetically diverse host populations select for parasites that can infect a broad range of host genotypes, while genetically homogeneous host populations select for specialist parasites. Prior studies have investigated the evolution of host range under temporal variation in the host population, predominantly in virus-cell culture systems. We complemented this prior body of work by testing spatial, rather than temporal, variation in host diversity, in a eukaryotic host-parasite system. Consistent with our hypothesis, selection in diverse host populations maintained a broad host range in parasite populations (Fig. 3). However, selection in homogeneous host populations led to the evolution of specialist parasites in only one of the two sympatric host genotypes (Fig. 2).

Selection in homogeneous and diverse host populations resulted in parasite populations with increased virulence against the host genotype N2 (Fig. 2A). In fact, parasites selected in diverse host populations increased in virulence to the same extent as parasites selected in homogeneous N2 populations, as indicated by the statistically indistinguishable mortality rate of hosts exposed to these different parasite populations. In the case of homogeneous N2-selected parasites, increased killing of N2 coincided with a contraction of host range, as indicated by a loss of killing against the novel host JU1395. In contrast, for diverse-selected parasites, increased killing of N2 was accomplished without a contraction of host range: diverse-selected parasite populations maintained genotypes with high killing ability and less between-lineage performance in a novel host environment (Fig. 3). Our results suggest that genetic diversity of the host population prevented the fixation of mutations that carry deleterious effects in alternate host environments due to antagonistic pleiotropy or relaxed selection. We do not know the extent of polymorphism in diverse-selected parasite lineages: they may be monomorphic generalists or polymorphic, with some genotypes specialized on N2. These findings more broadly suggest that parasite lineages from genetically diverse host populations will be more likely to emerge in novel host populations.

Selection in homogeneous and diverse host populations did not result in parasite populations with increased virulence against the host genotype LTM1 (Fig. 2B). This lack of adaptation corresponded to the maintenance of a broad host range: homogeneous LTM1-selected parasites showed no contraction of host range relative to control and diverse-selected parasites (Fig. 3). Our experimental evolution may have provided insufficient time for adaptation to LTM1. Initially high rates of killing by ancestral parasites could have slowed fixation of beneficial mutations if these are rarer in the LTM1 host environment than the N2 host environment. Consistent with this hypothesis, changes in virulence against LTM1 qualitatively matched the predicted changes, with slightly increased virulence of diverse-selected and homogeneous LTM1-selected parasites relative to control and homogeneous N2-selected parasites (Fig. 2). Compared to what we observed with adaptation to N2, these changes were smaller in magnitude with more variation around the mean. Additional generations of selection may produce stronger differentiation between treatments. We conclude that intrinsic differences between these host genotypes altered the rate at which specialization evolved and thereby the dynamics of emergence probability on novel host genotypes.

Prior studies of host range have similarly found that host range evolves differently according to the host species encountered. Fellous et al. (2014) selected populations of the spider mite *Tetranychus urticae* for growth on different plant species. They found variation in host range according to the plant species: selection on rosebay produced mite populations with consistent performance across plant species, while selection on tomato produced mite populations with more variable performance. In Turner et al. (2010), adaptation to homogeneous lines of human and canine cells reduced viral performance on novel cell lines for the human-adapted viruses but not the canine-adapted viruses. They argue that performance in canine cell lines is broadly correlated with performance in other host environments, such that a homogeneous host environment can indirectly select for parasites with broad host range. Our results suggest that the same argument may apply to host range evolution at the level of host genotype.

Much of our knowledge of host range evolution at the level of host genotype comes from studies of coevolving bacteria-phage systems. These studies provide indirect support for the idea that host populations that maintain genetic diversity select for parasites that can infect a broad range of host genotypes: relative to bacteria or phage evolution alone, coevolution maintains more diversity within bacterial host populations and selects for phages with broader host ranges (Poullain et al. 2008; Hall et al. 2010). We prevented coevolution in our study by preventing evolution of our host lines. Prior experimental coevolution studies in this system find that coevolution can maintain diversity in host populations (Morran et al. 2011). Based upon our results here, we then predict that, on average, parasite lineages passaged with coevolving host populations will maintain broader host ranges than parasites serially passaged with host populations that are homogeneous in space and time. Broadly, our results point to the significance of the local host population, in terms of both the identity and diversity of host genotypes present, in determining a parasite population’s potential to shift to new hosts.

## Author contributions

AKG conceived and directed the study, performed experimental evolution and assays, collected data, analyzed data, and wrote the manuscript. HSBB assisted in experimental evolution, performed assays, collected and analyzed data, and contributed to drafts of the manuscript. MJP, JL, RO and AK assisted in experimental evolution and assays. LTM conceived the study, provided guidance, collected data, and revised the manuscript.

## Acknowledgements

We are grateful to Dilys Osei for her contributions to experimental evolution. This work was supported in part by funds from the National Science Foundation (DEB-1750553) to LT Morran. AK Gibson was supported by the NIH IRACDA program Fellowships in Research and Science Teaching (FIRST) at Emory University (K12GM000680). Some strains were provided by the CGC, which is funded by the NIH Office of Research Infrastructure Programs (P40 OD0140440).

