## Supplemental Material and Tables for "The evolution of parasite host range in genetically diverse host populations"

**Supplement**

*Statistical Analyses*

Our analysis of variation in survival of LMT1 hosts produced a mixed-effects model with a singular fit. A singular fit can arise from overfitting. To address this problem, we treated parasite lineage as a fixed effect, as opposed to a random effect. We then re-fit the model as a negative bionomial regression using the glm.nb function in the package MASS (Venables and Ripley 2002). Treatment remained an insignificant predictor of variation in mortality of LTM1 (D = 4.37, df = 3, *p* = 0.224).

**Supplemental Tables**

**Table S1: Mean virulence of experimentally evolved parasites against sympatric host genotype N2**

|  | **No. survivors** | | **Mortality rate** | |  |
| --- | --- | --- | --- | --- | --- |
| **Parasite treatment** | *Mean* | *SE* | *Mean* | *SE* | **No. lines** |
| Ancestor | 87.25 |  | 0.823 |  | 1 |
| Control | 85.96 | 4.10 | 0.826 | 0.008 | 6 |
| 100% LTM1 | 100.38 | 10.22 | 0.797 | 0.021 | 6 |
| 50% | 62.96 | 3.67 | 0.873 | 0.007 | 6 |
| 100% N2 | 57.17 | 6.67 | 0.884 | 0.014 | 6 |
| *Total added* | *494* | *26* |  |  |  |

We calculated the mean values for Number of surviving worms and Mortality Rate by averaging the mean values obtained for the six parasite lineages per treatment, which were in turn obtained by averaging the values obtained for the four experimental replicates per lineage. Standard errors of the mean therefore reflect variation across the six parasite lineages (hence the lack of standard error for the ancestor, which is a single lineage). We calculated mortality rate as the number of living worms divided by the total added, which was calculated as the mean number of worms counted on 10 standard plates.

**Table S2: Mean virulence of experimentally evolved parasites against sympatric host genotype LTM1**

|  | **No. survivors** | | **Mortality rate** | |  |
| --- | --- | --- | --- | --- | --- |
| **Parasite treatment** | *Mean* | *SE* | *Mean* | *SE* | **No. lines** |
| Ancestor | 88.5 |  | 0.822 |  | 1 |
| Control | 80.54 | 2.68 | 0.838 | 0.005 | 6 |
| 100% LTM1 | 67.46 | 4.17 | 0.865 | 0.008 | 6 |
| 50% | 70.75 | 3.11 | 0.858 | 0.006 | 6 |
| 100% N2 | 88.17 | 7.48 | 0.823 | 0.015 | 6 |
| *Total added* | *498* | *25* |  |  |  |

Calculations are as indicated in Table S1.

**Table S3: Mean virulence of experimentally evolved parasites against novel host genotype JU1395**

|  | **No. survivors** | | **Mortality rate** | |  |
| --- | --- | --- | --- | --- | --- |
| **Parasite treatment** | *Mean* | *SE* | *Mean* | *SE* | **No. lines** |
| Ancestor | 11.75 |  | 0.956 |  | 1 |
| Control | 17.15 | 0.85 | 0.936 | 0.003 | 6 |
| 100% LTM1 | 20.23 | 1.91 | 0.925 | 0.007 | 6 |
| 50% | 16.01 | 0.83 | 0.941 | 0.003 | 6 |
| 100% N2 | 37.59 | 4.89 | 0.861 | 0.018 | 6 |
| *Total added* | *270* | *12* |  |  |  |

Calculations are as indicated in Table S1, except mean values for each parasite lineage were calculated from eight experimental replicates.

**Table S4: Variation in virulence of experimentally evolved parasite lineages against novel host genotype**

|  | **No. survivors** | | **Mortality rate** | |
| --- | --- | --- | --- | --- |
| **Parasite treatment** | *CV* | *95%CI* | *CV* | *95%CI* |
| *Control* | 0.299 | (0.155,0.582) | 0.020 | (0.010,0.042) |
| *100% LTM1* | 0.565 | (0.413,0.779) | 0.046 | (0.032,0.065) |
| *50%* | 0.312 | (0.219,0.552) | 0.020 | (0.014,0.035) |
| *100% N2* | 0.780 | (0.622,0.965) | 0.126 | (0.092,0.169) |

Coefficients of variation are calculated across the six parasite lineages within each treatment.
